# Ecological Insights from the Evolutionary History of Microbial Innovations

**DOI:** 10.1101/220939

**Authors:** Mario E. Muscarella, James P. O’Dwyer

## Abstract

Bacteria and Archaea represent the base of the evolutionary tree of life and contain the vast majority of phylogenetic and functional diversity. Because these organisms and their traits directly impact ecosystems and human health, a focus on functional traits has become increasingly common in microbial ecology. These trait-based approaches have the potential to link microbial communities and their ecological function. But an open question is how, why, and in what order microorganisms acquired the traits we observe in the present day. To address this, we reconstructed the evolutionary history of microbial traits using genomic data to understand the evolution, selective advantage, and similarity of traits in extant organisms and provide insights into the composition of genomes and communities. We used the geological timeline and physiological expectations to provide independent evidence in support of this evolutionary history. Using this reconstructed evolutionary history, we explored hypotheses related to the composition of genomes. We showed that gene transition rates can be used to make predictions about the size and type of genes in a genome: generalist genomes comprise many evolutionarily labile genes while specialist genomes comprise more highly conserved functional genes. These findings suggest that generalist organisms do not build up and hoard an array of functions, but rather tend to experiment with functions related to environmental sensing, transport, and complex resource degradation. Our results provide a framework for understanding the evolutionary history of extant microorganisms, the origin and maintenanceof traits, and linking evolutionary relatedness and ecological function.

---

Global estimates predict upwards of 10^12^ bacterial and archaeal species (1, 2); however, the origin and maintenance of traits that define their phenotypes remains largely unknown. Trait evolutionary history, which captures these processes, is the foundation describing differences between species and links evolutionary relatedness and ecological function (3). The true evolutionary history is unknown, but observed traits provide a window into evolutionary processes through their distributions across phylogenic trees. To better understand these processes we can model the trait distributions. A simple model would be that traits are conserved and therefore maintained by descendants. As such, taxonomy and/or phylogeny could be used as a trait proxy because specific traits would be conserved within distinct groups (4, 5). Here, we refer to this as the *conservation framework*. This framework is common in microbial ecology (6). It has been used to predict traits (4), identify functional groups (7), and is often used to describing differences between communities based on indicator taxa. A more complex model would be that traits can be gained through innovation but also lost by descendants due to changes in selection and environmental context (8). In this model, the dynamics of trait gain and loss can be represented as a stochastic process (9, 10). Here, we refer to this as the *gain-loss framework*. This framework is a common way to infer ancestral states (11) and to predict traits for unknown organisms (12). While lacking nuance related to specific ecological and evolutionary mechanisms, both frameworks can describe the emergent trait distribution patterns observed with extant organisms.

These frameworks differ in the trait history they predict, but the differences provide an opportunity to understand trait innovation and selection. For example, the conservation framework assumes that closely related organisms have similar traits and functions (13, 14). This assumption leverages the hierarchical evolutionary ancestry of organisms and the idea that related organisms should resemble each other (15, 16). In contrast, the gain-loss framework assumes that both trait gain and loss are common evolutionary mechanisms that can confer fitness advantages (17–19). While there has not been a wide-scale quantitative comparison of these frameworks, there is no reason to expect one framework across all traits. For example, while we know there is a basic set of genes required for cellular function (20), high levels of genome reduction have been documented (21). As such, it may be more informative to compare predictions across traits. When the frameworks agree, it would suggest that the loss rate is small and therefore trait loss by descendants may be strongly selected against. This would be expected for genes related to cellular division and central metabolism. In contrast, if the frameworks disagree then it would suggest that trait loss is high and therefore traits can be easily lost by descendants. Traits with high gain-loss rates (*i.e.*, evolutionarily labile) could provide mechanisms for evolutionary diversification between related species.

Trait evolutionary history provides unique ecological insights and can be used to understand the composition of genes in a genome and species in an ecosystem. For example, genome size reflects the long term interaction between gene gain and loss (22). As such, understanding the types of genes with low loss rates (*i.e.*, evolutionary conserved) and those with high gain and loss rates (*i.e.* evolutionary labile) will provide a new way to probe the differences between generalists with large genomes and specialist organisms with (typically) smaller genomes. Likewise, evolutionary history provides a unique context for community structure–function relationships. Across the tree of life, ecological functions are clustered at different evolutionary depths (6), and functions observed at recent nodes are often highly dispersed throughout the tree of life (13). The evolutionary history of genes provides the quantitative framework to explain these patterns. As such, it will help determine which ecological and physiological functions are evolutionarily constrained and which are dispersed, and therefore determine which functions we expect to be performed conserved phylogenetic clades versus redundant groups.

In this study, we use 3179 bacterial and archaeal genomes to explore trait innovation based on individual genes. In total, 2950 orthologous genes were associated with the genome collection. Genes represent the raw genetic information underlying traits; therefore, while not phenotypic traits themselves, genes can be used to understand trait distributions. Using these genomes, we describe microbial innovations using the frameworks outlined above and illustrated in Fig. 1. Specifically, the conservation framework identifies phylogenetic nodes where 90% of the descendent genomes contain a gene of interest (13). In contrast, the gain-loss framework estimates the nodes where traits first arose using a two-state Markov process, allowing for traits to be both acquired and lost over time (9). Using these predictions, we compare the inferred innovation dates across genes to understand the evolutionary history and to make predictions regarding trait selection and genome composition.

**Fig. 1.**
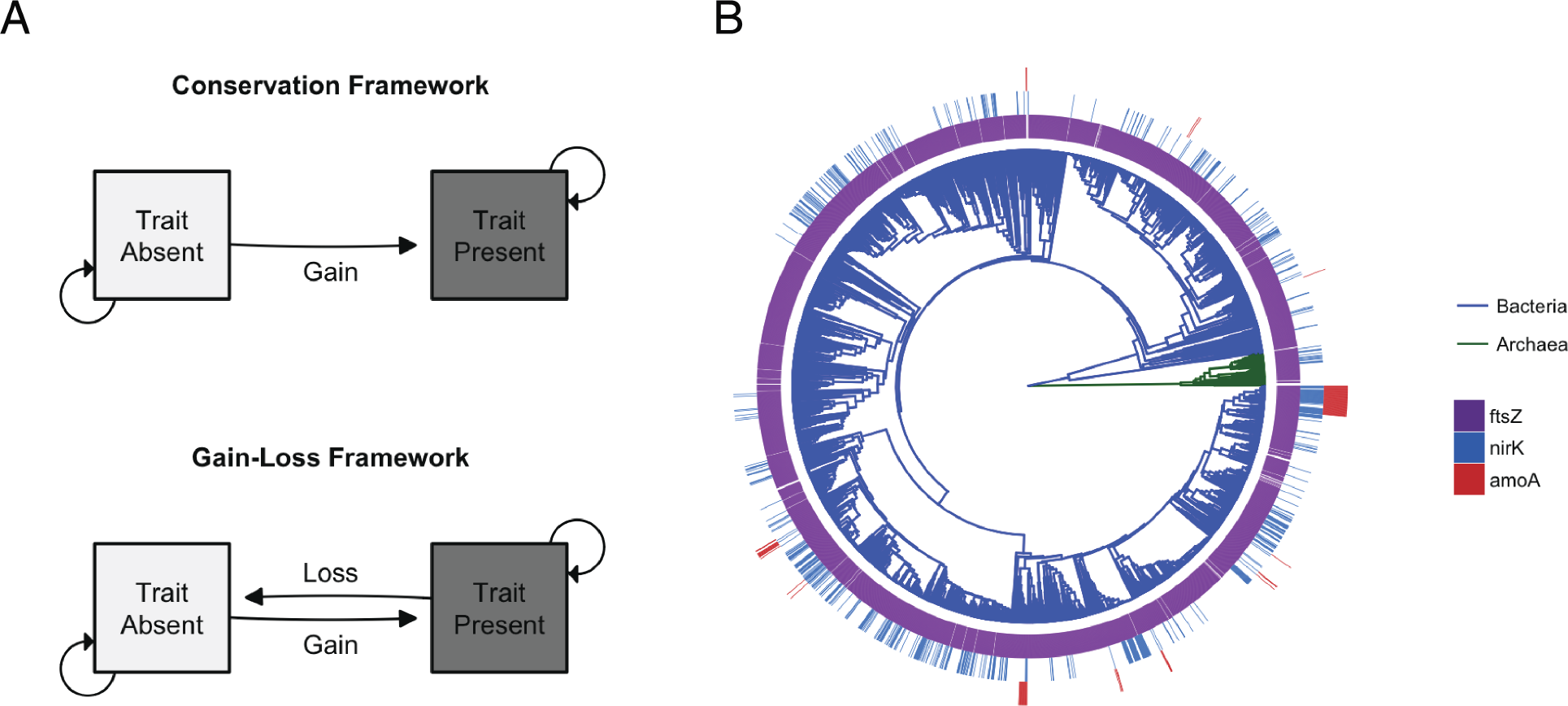
Trait innovations in the microbial tree of life. **A** The two frameworks for trait innovation. Under the conservation framework traits are gained but not lost. Absent traits are maintained with probability of *A* or gained at probability 1 - *A*. Once a trait is gained, it is maintained by descendants. Under the gain-loss framework traits are gained and lost. Absent traits are maintained with probability *A* or gained at probability 1 - *A*. Once a trait is gained, it is maintained at probability *B* or lost at probability 1 - *B*. **B** Phylogenetic tree of bacteria (n = 3108) and archaea (n = 71). The tree is based on the 16S rRNA sequences. Example genes are plotted at the tips. Some genes (e.g., *ftsZ* : cell division ring) are found in almost all genomes. Other genes (e.g., *nirK*) are abundant on the tree, but highly dispersed across taxa. Finally, some genes (e.g., *amoA*) are found in only a few groups. The phylogenetic tree is available in Newick format in Supplementary Dataset 1 and the gene occurrence table is available in Supplementary Dataset 2.

## Results and Discussion

Across genes, we find that predictions under the gain-loss framework are more ancestral than those for the conservation framework (Fig. 2A). However, there is a large range in the discrepancy (Fig. 2B). This suggests that the frameworks are part of a spectrum describing gene evolutionary history. At one end, genes are essential and highly conserved after they originate. On the other end, genes are non-essential and can be easily lost by descendants. Our results suggest that the essential to non-essential gradient is related to inferred gene loss rates (Fig. 2C). To test whether we can generate accurate inferences for gain and loss on a similarly sized tree, we simulated an idealized system where the true dynamics are controlled by a Markov process (See Supplemental). In this simulation, we used a tree with similar size and age, and a range of transition parameters which were similar to the inferred rates. In addition, our simulation tested the prediction accuracy across the range of transition rates we observed. Our simulation confirms that the observed discrepancy can be linked to high loss rates, and it provides further evidence to suggest that the conservation framework is a special case within the gain-loss framework where loss rates are extremely low. Our findings also suggest that loss rates which lead to discrepancy are common across most genes (85%), which is consistent with studies showing high levels of genome reduction throughout evolutionary time (21). Therefore, our findings suggest that the inferred gene loss rate is a quantitative measure of gene evolutionary lability (*i.e.*, easily gained or lost), and can be used as a proxy for how essential a gene is for cellular function.

**Fig. 2.**
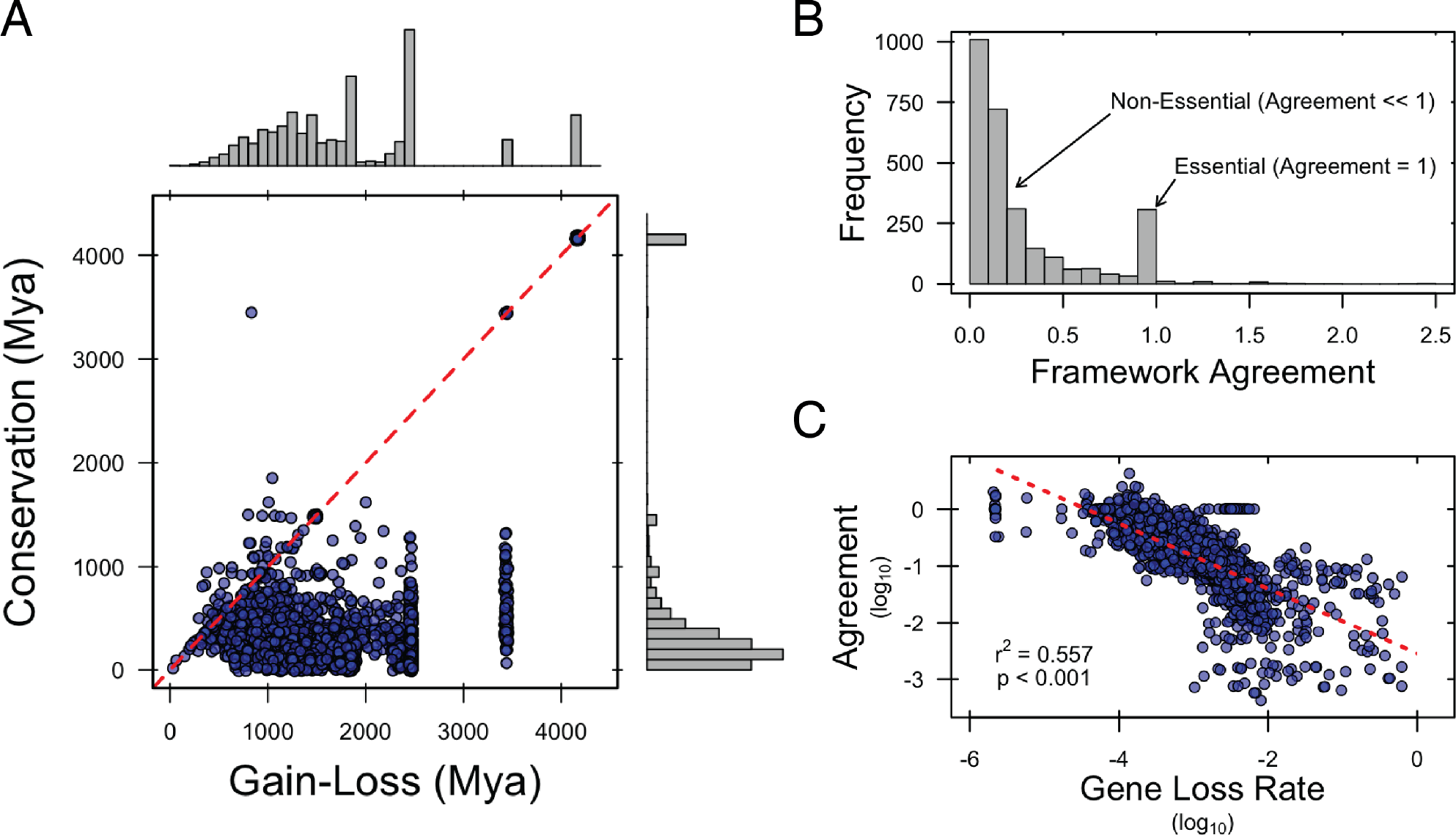
Innovation predictions under the gain-loss and conservation frameworks. **A: Comparison between frameworks.** Across genes, we find that gain-loss (GLF) predictions are more ancestral than those for conservation (CF). The CF identifies two major groups of genes. The first group is at the tree root contains 192 genes, including *ftsZ* (cell division), *dnaA* (DNA replication), *GPI* (glycolosis), as well as other genes for cell growth and division, translation, and oxidative phosphorylation. The second group includes 2666 genes at more recent nodes (~ 222 Mya). Most (95%) are associated with metabolism. In contrast, the GLF predicts more variation. First, a group of 229 genes is predicted at root, with a 99% overlap with the CF. We find peaks of innovation around 3400, 2400, and 1800 Mya. Finally, we find 1577 recent innovations (~ 1200 Mya). Similar to the CF, most are associated with metabolism. **B: Agreement between the frameworks.** The agreement between the frameworks was calculated as the CF estimate divided by GLF estimate. If the frameworks agree, then the agreement would be equal to 1. Across genes, we find 290 genes with agreement between the frameworks. This suggests that once these genes evolve, they are maintained by descendants. However, about 2500 genes have an agreement below 1. This suggests that these genes are not maintained after they originate. **C: Lower agreement is associated with higher gene loss rate.** Loss rates and agreement were log_10_ transformed and a linear regression model was used to determine the relationship between loss rate and agreement. A significant negative relationship was found (F_1,2796_ = 3388, p < 0.001), suggesting that the difference in agreement between the frameworks is related to the loss rate inferred by ancestral state reconstruction.

## Innovation Origins Match Physiological and Geological Expectations

To explore predictions regarding the origin of innovations, we compared the predictions for genes within broad pathways. Pathway characteristics (*e.g.*, required for cell division) and Earth’s geological history can be used to provide independent lines of evidence with which to test our inferences. For example, while some cellular processes are required by all organisms and thus show strong agreement at the base of the tree (*e.g.*, *fitZ* – encodes the cell division ring) other processes range from group specific (e.g., *mutH* – encodes a sequence specific endonuclease) to evolutionarily labile (*e.g.*, *solR* – a quorum-sensing system regulator) (Fig. 3A). Likewise, we can leverage Earth’s geological history as independent evidence. For example, current estimates suggest that life originated during the Hadean (*>* 4000 Mya) (23), and that chemoautotrophic organisms dominated during the Archaean (4000 – 2500 Mya) (24). During the Archaean anoxygenic photosynthesis originated (25) and towards the end of the eon oxygenic photosynthesis originated thus ensuing the Great Oxidation Event (24). This series of geological events qualitatively supports the ordering of our predictions (Fig. 3B). Furthermore, the discrepancy between the frameworks suggests that while oxygenic photosynthesis is maintained by a specific group (*i.e.*, cyanobacteria), anoxygenic photosynthesis is highly labile and often lost by descendants. Together, our results qualitatively recapitulate the evolutionary history of microbial traits and further suggest that the differences between frameworks is a signal of evolutionary lability.

**Fig. 3.**
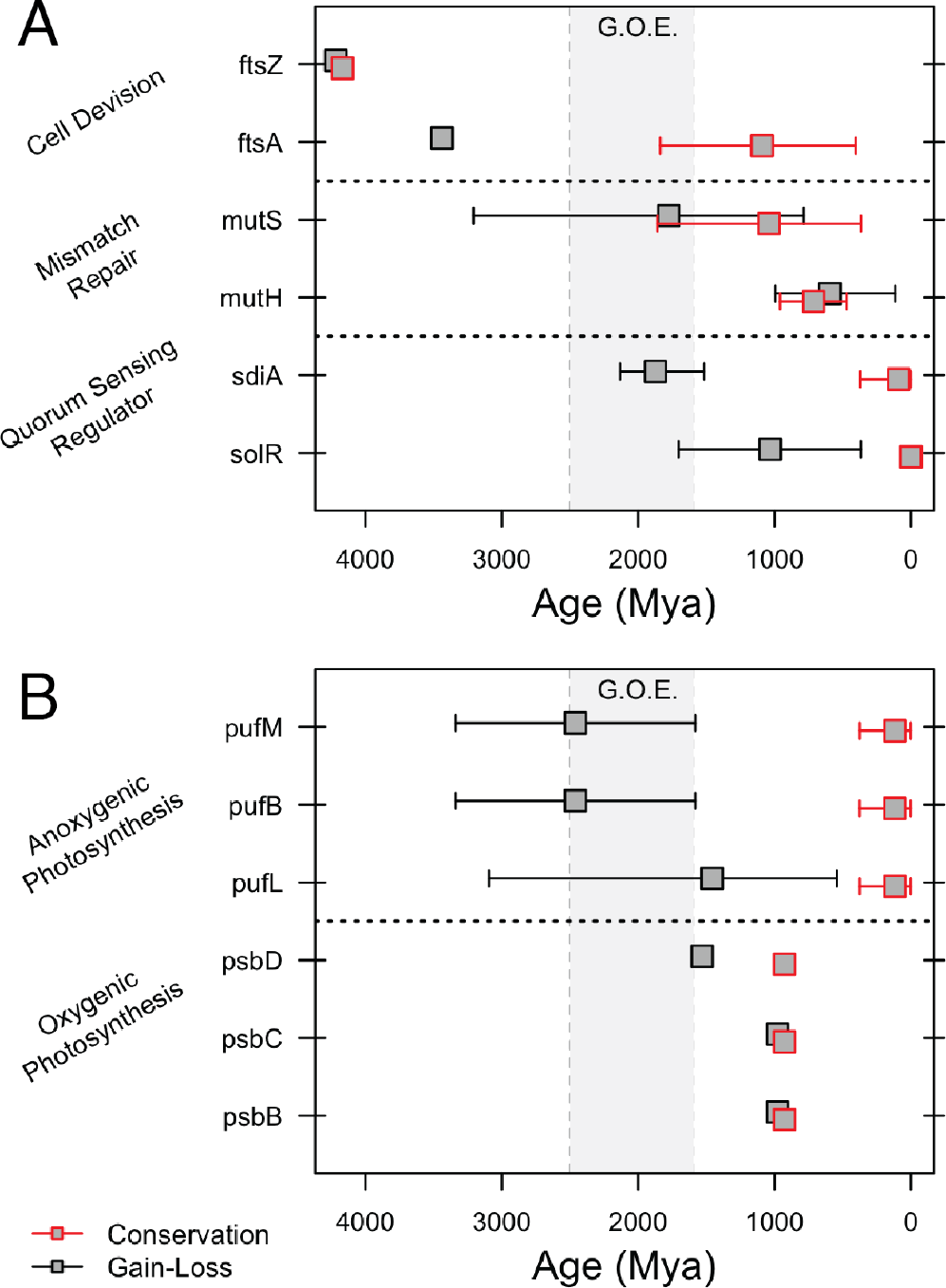
Predicted innovation dates for genes in various pathways. Example gene innovations predicted under the conservation (*red boxes*) and gain-loss (*gray boxes*) frameworks are shown for cellular processes (**A**) and photosynthesis (**B**). For some genes, there is a strong agreement between the two frameworks and the predictions overlap (*e.g.*, *ftsZ*). For other genes, there is little agreement between the frameworks. For example, genes related to anoxygenic photosynthesis show a much earlier origin under the gain-loss framework. In addition, predictions under the gain-loss framework are in better agreement with geological evidence for the appearance of specific metabolic processes. For example, the Great Oxidation Event is predicted to have taken place between 1800 and 2500 Mya (25). Before this event, oxygen was a trace element in the atmosphere and anaerobic processes dominated. In general, the predictions from the gain-loss framework qualitatively recapitulate the ordering of these predictions.

### Gene Loss Rate is a Measure of Evolutionary Lability

To explore the evolutionary lability predictions, we compared the predictions for specific genes. For example, there are about 290 genes with a strong agreement between the frameworks (Fig. 2B). Genes predicted to originate at the base of the tree are involved in cellular processes such as growth, information processing, and central metabolism which are required by all organisms. Likewise, genes involved with oxygenic photosynthesis are known to be necessary for specific groups of microorganisms (*i.e.*, cyanobacteria). We would expect the loss of these genes is strongly selected against in most circumstances, and indeed they have low loss rates and strong agreement between the frameworks. In contrast, there are *>*2500 genes with disagreement between the frameworks. The majority of these genes (*>*70%) are involved with cellular metabolism and 17% are involved with environmental information processing. While these genes are not needed for core cellular functions (*e.g.*, DNA replication, cell division), they are used to acquire resources and contend with environmental variation. We expect these genes to be more evolutionarily labile, and we find that they have high loss rates and low agreement between the frameworks. Therefore, these genes may represent ways organisms experiment with new traits and diversify into new ecological niches.

### Evolutionary History is Linked to Genome Evolution and Ecology

Documenting the evolutionary history of traits allows us to gain insights into the ecology and evolution of organisms. Here, we focus on how origin and maintenance of traits describes the development of generalist and specialist organisms. The number of distinct metabolic pathways can be used to characterize the generalism to specialism gradient (26), and this number could increase by either accumulating of genes over time or constantly acquiring and purging genes. We refer to the first scenario as the *gene hoarding hypothesis* which would be supported if the average gene loss rate in a genome was constant with genome size. We refer to the second scenario as the *gene experimentation hypothesis* which would be supported if the average loss rate in a genome increased with genome size. Across genomes, we find a strong positive linear relationship between the estimated gene loss rates and genome size (Fig. 4). Furthermore, we found that genes in the core genome (*i.e.*, genes shared across organisms) had lower loss rates than genes indicative of large genomes (Fig. 5A). These findings suggest that larger genomes contain genes which, on average, have higher loss rates (Fig. 2C). Therefore, if genome size is a signature of the generalist–specialist gradient, then generalists contain more evolutionarily labile genes and specialists contain more physiological essential genes. Furthermore, while generalists maintain more genes, they are not hoarding traits but rather experimenting with new ecological functions. These findings suggest that generalists are undergoing ecological diversification through a process of trait experimentation while specialists are relying on conserved functional genes.

**Fig. 4.**
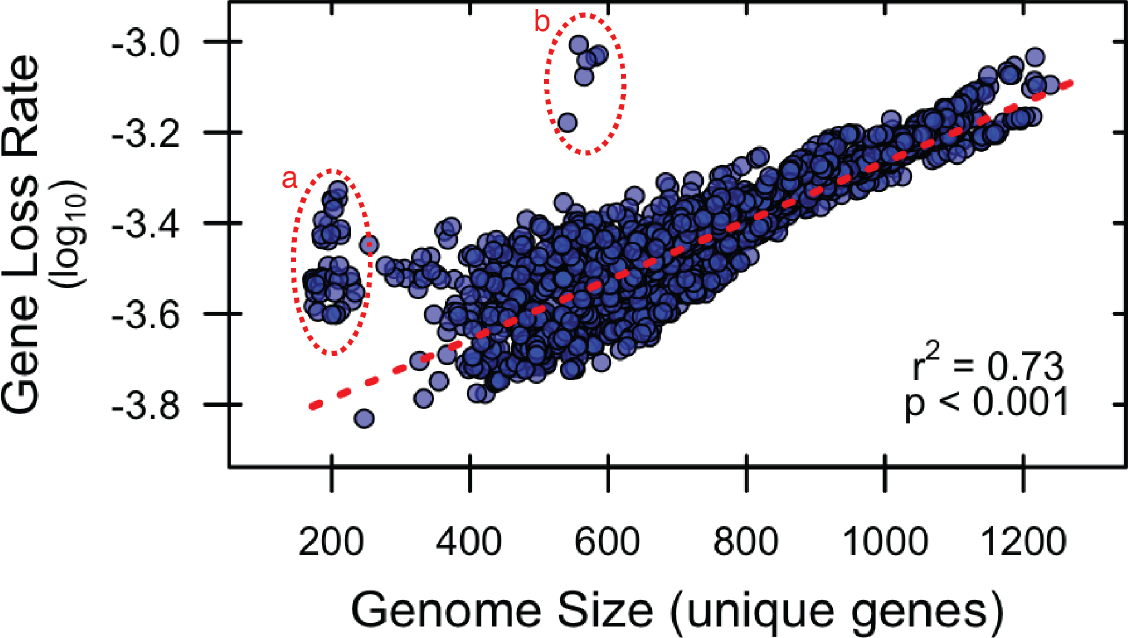
Larger genomes contain genes with a higher gene loss rate. There is a linear relationship between the number of unique annotated genes in a genome and the median gene loss rate of the genes contained in the genome. The loss rate of each gene is the maximum likelihood estimate for the gene switching rate based on the ancestral state reconstruction. Loss rates were log_10_ transformed and a linear regression model was used to determine the relationship between genome size and loss rate. A significant positive relationship was found (F_1,3123_ = 8657, p *<* 0.001). Note: *a* – endosymbiont genomes which have lost some essential functions, *b* – Caulobacter genomes which have unique cellular division process and have lost the common pathways.

**Fig. 5.**
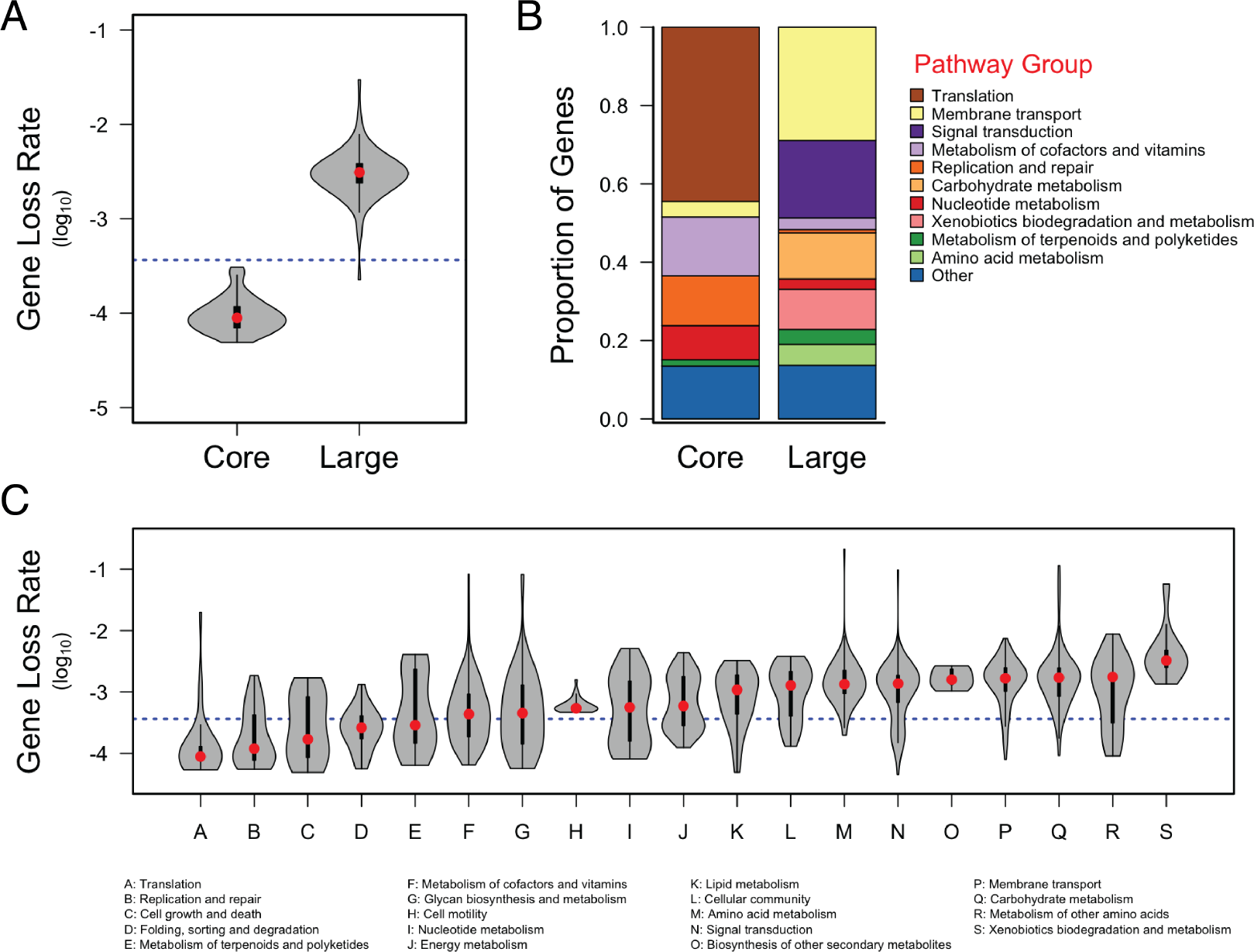
Genome loss rates and composition. **A** Loss rates for genes characteristic of the core genome and large genomes. Violin plots show the kernel density for each genome group. The core genome was defined as those genes found in greater than 85% of genomes. The large genome was defined as those genes that are statistically more likely to be found in genomes which were 2 standard deviations larger than average. The blue line represents the median loss rate across all genes. The red dots represent the median loss rate for core and large genome genes. **B** KEGG pathway groupings, based on KEGG hierarchy level B, for the genes in the Core and Large genome groups. Stacked barplots represent the proportion of genes in each group. **C** Gene loss rates for all genes based on KEGG pathway groupings. Violin plots show the kernel density of each pathway group. The blue line represents the median loss rate across all genes. The red dots represent the median loss rate for each pathway group.

Trait evolutionary history provides insights into genome composition and ecological functioning. To explore genome composition, we grouped genes based on the KEGG ontology hierarchy. We found major composition shifts between core and large genome genes (Fig. 5B). As expected, the core genome contains functions related to growth (translation), reproduction (replication and repair), and cellular requirements (vitamins and cofactors). In contrast, genes unique to large genomes belong to pathways for membrane transport, regulation (signal transduction), carbohydrate metabolism, and xenobiotic biodegradation. These findings suggest that large genomes acquire functions allowing them to use novel resources and sense environmental conditions, and they demonstrate that specific ecological functions drive the pattern between genome composition and gene loss rate. To explore ecological functions, we compared genes loss rates based on their KEGG pathway groupings. We found that pathway groups with loss rates below average include functions such as translation, replication and repair, cell growth and death. In contrast, pathway groups with loss rate above average include membrane transport, carbohydrate metabolism, and xenobiotic degradation. While most groupings had unimodal loss rate distributions (*e.g.*, Translation, Membrane Transport), some groupings have bimodal distributions (*e.g.* Energy Metabolism). In these cases, we looked at the specific genes along the distribution and found distinct genes in each mode. For example, the lower mode in energy metabolism includes genes related to ATP synthesis and methane metabolism while the upper mode includes genes related to nitrogen and sulfur metabolism (Fig S5). This higher resolution analysis also explains the tails observed for some distributions. For example, the lower tail associated with signal transduction includes genes involved in cellular nitrogen and phosphorus starvation responses (Fig S6). Together, our results demonstrate the link between evolutionary loss rates and ecological function, and we expect functions with higher loss rates to be dispersed and performed by functionally redundant clades.

## Conclusions

In this study, we inferred and interpreted the evolutionary history of microbial traits. We used two frameworks which differ in their assumptions, and we demonstrated that the framework which included loss met the physiological and geological expectations for microbial evolutionary history. We found that organisms with large genomes contain genes with higher loss rates and these genes belong to specific pathways. In addition, we found that when genes are grouped based on their physiological and ecological functions they have similar loss rates. Functions related to cellular growth and reproduction have the lowest loss rates and functions related to sensing environmental conditions and use complex resources have the highest loss rates. Given the relationship between loss rate and framework agreement, our findings suggest that the conservation framework would not be an accurate predictor of the evolutionary history for most innovations. As such, the conservation framework is best when used as a heuristic to group extant organisms without direct evolutionary implications or assumptions about trait loss. In contrast, while the gain-loss framework lacks mechanisms such as horizontal gene transfer, varying rates, and the non-independence of trait gain and loss, it qualitatively recapitulates the evolutionary history of major metabolic processes. In addition, while some traits may confer changes in diversification rates (27), it is implausible that these would be sustained indefinitely. Together, our results provide a tractable framework for understanding microbial evolutionary history and describing the evolutionary underpinnings for ecological differences between species.

## Materials and Methods

Genomes were downloaded from the Joint Genome Institute (JGI) Integrated Microbial Genomes (IMG) data warehouse. Briefly, we downloaded the full list of publicly available bacterial and archaeal genomes (accessed April 2017, Supplemental Table 1). Using the JGI project ID, we searched the JGI project status page for the database name and the names of any larger project databases. Using the JGI Genome Portal API, we then searched for project archives and downloaded the most recent complete archive. We extracted each archive and only included those representing genomes with KEGG annotations and high quality 16S rRNA gene sequences (*>* 1200 bp). The 16S rRNA gene sequences were identified by searching the annotation file (*.gff) for entries based on the following search criteria: ‘rRNA.*product=16S‘. We parsed the entries for annotated gene IDs and searched the associated sequence file (*.fna) for sequence entries. The KEGG KO annotations were found in the KO table file associated with each genome (*.ko.tab.txt). We used KEGG annotations because they are hierarchically organized into pathways and modules (28). In total, 3179 genomes met the criteria to be included in our study and together these genomes contained 2950 orthologous genes based on KEGG annotations. Database parsing and downloading was done using a custom automated script which retrieved the required information and downloaded the genomes using the JGI API. Genome parsing was using custom bash scripts and code implemented in R (29).

Using the 16S rRNA genes, we created a representative phylogenetic tree. We aligned the 16S rRNA genes based on the GreenGenes reference phylogeny (v. 13.8.99) using mothur (30). We only included sequences which aligned to the reference. We used FastTree to generate a phylogenetic tree assuming the general time reversible model of nucleotide evolution (31). We applied midpoint rooting to our tree and used treePL to estimate divergence times (32). We standardized the tree by setting the root at 4000 *±* 200 Mya (23), estimates at this date should be regarded as evolving prior to the bacterial–archaeal divergence. To prevent bias when comparing predictions to geological events, we did not internally calibrate our tree. Therefore, the dates inferred in this study should only be used as qualitative estimates. To check the accuracy of our tree reconstruction, we compared taxonomic assignments with tree topology. The phylogenetic tree is available in Newick format in Supplementary Dataset 1.

We used KEGG annotations to infer the evolutionary history of traits. While genes do not represent phenotypic traits, they represent the genomic underpinning for traits and provide a standardized method to compare organisms. We treated genes as discrete traits based on presence-absence. We then inferred the evolutionary history using our proposed frameworks (Fig. 1). Under the conservation framework, we identified the nodes where 90% of the downstream genomes contained the gene of interest (13). Under the gain-loss framework, we fit a continuous time discrete two-state Markov model to the observed trait states using maximum likelihood estimation (11). This is the commonly used Mk2 model and we used the joint likelihood for maximum likelihood estimation. Other models of ancestral state reconstruction exist, including those which allow for state dependent diversification rates (27), but we assumed that changes in diversification rate would not be sustained at the evolutionary timescale of our tree. Using the inferred rate parameters, we estimated the probability of trait states at each node using the posterior probabilities at each internal node. We identified the trait *innovation* as the first node at which the trait most likely went from absent to present in a lineage using a posterior threshold of 0.5. Both methods were implemented in R using code adapted from the custom *ConsenTrait* function (13) and the *fitMK* function from the *phytools* R package (33) in addition to custom scripts. The gene occurrence table is available in Supplementary Dataset 2.

We used the inferred loss rates to made inferences based on genome size and gene pathway. To determine if there was a relationship between genome size and loss rate we used linear regression. We used the median loss rate across all genes in each genome, and we used the number of unique homologous genes as the measure of genome size. To determine how the composition of genes relates to the ecological function, we first defined the core and large genome. The core genome was defined as those genes found in *>* 85% of genomes. The large genome was defined as those genes unique to large genomes. To determine which genes were unique to large genomes we used logistic regression. First, we z-transformed genome size so that we could define large genomes as those at least two standard deviations larger than the average genome size. We then fit a logistic regression using gene presence as the response variable and transformed genome size as the predictor. The genes unique to the large genome were defined as those with significant logistic regressions and a 50% chance of the gene being present in a large genome. To group genes based on physiological and ecological functions, we used the KEGG Ontology hierarchy level B. We refer to these groups as “*pathway groups*”.

## Acknowledgments

We thank K. Webster, S. Kembel, and J. Martiny for critical feedback on an earlier version of this manuscript. JOD acknowledges the Simons Foundation Grant #376199, McDonnell Foundation Grant #220020439, and NSF Grant #DEB1557192.

